# Women’s estimated HIV infections from sex in trials of pre-exposure prophylaxis in Africa: Implications for HIV prevention strategies

**DOI:** 10.1101/146530

**Authors:** David Gisselquist

## Abstract

**Introduction:** During 2004-15, nine randomized controlled trials (RCT) for HIV prevention tested pre-exposure prophylaxis (PrEP) with oral drugs, vaginal gels, or vaginal rings among more than 17,000 women in Africa.

**Methods:** This study uses information from the nine RCTs to estimate the proportions of HIV from sexual and bloodborne risks, to consider reasons for success or failure with oral PrEP, and to consider risks with vaginal PrEP.

**Results:** Estimating from women’s reported frequencies of unprotected coital acts in six RCTs, only a minority of women’s infections came from sex. Oral PrEP may have succeeded in at least one trial by reducing infections from both bloodborne and sexual risks. Oral PrEP may have failed in several trials, at least in part, because some women used oral PrEP when they had sexual risks rather than daily as advised. Relatively high incidence with PrEP vaginal gels and rings vs. oral placebo suggests vaginal PrEP had little impact at best and may have been harmful.

**Discussion:** Evidence from this and other studies challenges the common belief most HIV in Africa comes from sex. This challenge has implications for HIV prevention strategies, including: warning about bloodborne risks; and reconsidering PrEP for young women.

## Introduction

Outside Africa, less than 0.3% of adults are HIV-positive, more than twice as many men as women, and infections concentrate in men who have sex with men (MSM) and injection drug users (IDU).[1] In Africa women account for more infections than men, and their lifetime risk exceeds 25% in much of southern Africa.[1,2]

Unlike many diseases such as colds and malaria which spread through the air, casual contact, or common insect vectors, HIV transmits through easily recognized events: anal and vaginal sex and skin-piercing procedures. Thus, a core strategy for HIV prevention throughout the world has been to educate people to recognize and control risks. From this perspective, women’s high prevalence in parts of Africa shows they have failed to do so. Common explanations for this failure are that they have been unwilling or unable to limit sexual risks. Although such explanations agree with anecdotes (eg, some men refuse to use condoms), they do not fit survey data showing women in Africa with similar or less risky sexual behavior compared to women in developed countries.[3,4]

Another possible explanation for women’s failure to control their risks is that public health messages have not warned them about all important risks, ie, bloodborne as well as sexual risks. This explanation shifts the blame for failure to control risks from women with HIV infection to designers and disseminators of incomplete public health messages.

Researchers have tried for decades to find strategies to protect women in Africa from HIV. During 1987 to September 2011, 38 randomized controlled trials (RCTs) of vaginal gels, treating sexually transmitted infections (STI), and other interventions to reduce HIV incidence among women (or men and women) in Africa reported results.[5] Successes were rare. One of nine RCTs testing STI treatment[6] reported significantly lower incidence in the intervention vs. control arm; that trial overlapped an injection safety initiative.[7] Aside from that suspect success, the only RCTs to report significantly lower incidence among women (or men and women) in intervention vs. control arms were: a trial of treatment as prevention among discordant couples[8]; and three trials of pre-exposure prophylaxis (PrEP).[9-11] One other RCT of PrEP among women had much lower incidence in the intervention arm, but the result was not significant[12]; while another RCT of PrEP showed no impact.[13]

During October 2011 through 2016, four more RCTs of PrEP among women in Africa reported results. Considered together, during 2004-2015, nine RCTs of oral and vaginal PrEP in Africa followed more than 17,000 women for more than 22,000 person-years (PYs), observing 980 incident infections (Table 1). This paper uses evidence from these nine RCTs to estimate the proportions of women’s HIV from sexual risks. These proportions, along with other evidence from these trials, can inform the design of more effective HIV prevention activities.

**Table 1:** Women’s participation in RCTs for PrEP in Africa

| Study [reference] | Location | study years | Number followed | Follow-up PYs | Incident infections |
| --- | --- | --- | --- | --- | --- |
| TOTAL |  | 2004-15 | 17,708 | 22,501 | 980 |
| Phase II TDF, NCT00122486[12] | GH, CM, NGA | 2004-06 | 859 | 474 | 8 |
| CAPRISA 004[9] | SA | 2007-10 | 889 | 1,341 | 98 |
| TFD2, NCT00448669[14] | BT | 2007-10 | 548* | 714* | 33 |
| Partners PrEP, NCT00557245[15] | KY, UG | 2008-11 | 1,794* | 2,975* | 45 |
| FEM-PrEP, NCT00625404[16] | KY, SA, TZ | 2009-11 | 2,056 | 1,407 | 68 |
| VOICE, NCT00706579[17] | SA, UG, ZI | 2009-12 | 4,969 | 5,469 | 304 |
| FACTS 001, NCT01386294[18] | SA | 2011-14 | 2,029 | 3,036 | 123 |
| Ring Study, NCT01539226[19,20] | SA, UG | 2012-15 | 1,950 | 2,805 | 133 |
| ASPIRE, NCT01617096[21] | MW, SA, UG, ZI | 2012-15 | 2,614 | 4,280 | 168 |

Abbreviations: PY: person-years. TDF: tenofovir disproxil fumarate. BT: Botswana. CM: Cameroon. GH: Ghana. KY: Kenya. MW: Malawi. NGA: Nigeria; SA: South Africa. TZ: Tanzania. UG: Uganda. ZI: Zimbabwe.
\* These data are estimated as follows: TFD2 followed 1,200 men and women for 1,563 PYs; numbers and PYs of women followed are estimated from the report women were 45.7% of participants randomized. Partners PrEP reports 4,722 couples with at least one post-randomization HIV test and 7,830 PYs of follow-up; numbers and PYs of women followed are estimated from the report men were 62% of HIV-negative partners in couples followed.

## Methods

Recent reviews[5,22-24] identify nine RCTs of PrEP among women (or men and women) in Africa. For these RCTs, I use information in refereed medical journals, conference abstracts, and other documents: to estimate women’s HIV incidence from reported sexual behavior (for six trials disclosing women’s reported frequency of unprotected coital acts); to consider women’s awareness of sexual risk as a factor in their decisions to adhere to daily oral PrEP (for five trials of daily oral PREP); and to consider risks with PrEP gels and rings.

My search for relevant information has been guided by questions, eg: What did women report about sexual behavior? Why did women not take daily pills? Information from most RCTs is distributed across multiple documents. Some reports are pending, and I may have missed some published information. However, consistency among what I have found suggests missing evidence would have little impact on analyses in this paper.

## Results: Estimated HIV incidence from sexual and bloodborne risks

During monthly follow-up visits, all studies tested women for HIV and asked about sexual behavior (except FEM-PrEP, which asked quarterly). Six of nine RCTs (Table 2) disclosed sufficient information to estimate infections from sex (reported coital frequency; reported condom use over time or at last coital act). Three of these six studies disclosed frequency of unprotected coital acts during follow-up (FEM-PrEP did so for the last follow-up visit only); in all three studies, frequency of unprotected coital acts fell from baseline to follow-up. The other three of these six studies (Table 2) disclosed frequency of unprotected coital acts reported at baseline but not follow-up. Aside from these six, the remaining three of nine RCTs (TDF2, Partners PrEP, and FACTS 001) have not reported coital frequency and condom use at baseline or follow-up (as of May 2017); it’s not clear if the information they collected on sexual behavior included those details.[14,15,25]

**Table 2:** Women’s expected HIV incidence from reported sexual behavior in six RCTs

| Study, sources | Sexual behavior reported at follow-up (F) or baseline (B) |  | Partners with HIV (from annex Table 1) | Expected infections from sex |  | Total number of incident infections (from Table 2) |
| --- | --- | --- | --- | --- | --- | --- |
|  | Number of coital acts | % w/o condoms |  | infections/100 PYs (= coital acts/year x % w/o condoms x % of partners with HIV x 0.001) | Number (= incidence rate x PYs of follow-up [from Table 2]) |  |
| Phase II TDF [12] | 15/week (F) | 8% (F) | 3.3% | 0.2 (= 780/year x 8% x 3.3% x 0.001) | 1 (= 0.002 x 474) | 8 |
| CAPRISA 004 [9] | 5.0/month (F) | 20% (F) | 19.5% | 0.2 (= 60/year x 20% x 19.5% x 0.001) | 3 (= 0.003 x 1,341) | 98 |
| FEM-PrEP [16] | 3.41/week (F,B*) | 49% (F,B*) | 14.1% | 1.1 (= 177/year x 49% x 14.1% x 0.001) | 17 (= 0.012 x 1,407) | 68 |
| VOICE [17] | 2.5/week (B) | 15% (B) | 17.7% | 0.3 (= 130/year x 15% x 17.7% x 0.001) | 19 (= 0.003 x 5,469) | 304 |
| Ring Study [19,20] | 8.0/month (B†) | 38% (B) | 17.8% | 0.6 (= 96/year x 38% x 17.8% x 0.001) | 18 (= 0.006 x 2,805) | 133 |
| ASPIRE [21] | 26.5 last 3 months (B) | 43% (B) | 15.9% | 0.7 (= 106/year x 43% x 15.9% x 0.001) | 31 (= 0.007 x 4,280) | 168 |
| Median or total | Median: 118/year | Median: 22% | Median: 16.8% | Median: 0.5 per 100 PYs | Total: 89 | Total: 779 |

Abbreviations: PY: person-years. w/o: without. TDF: tenofovir disoproxil fumarate
\* These data average sexual behavior reported at baseline and at the last follow-up: 3.7 coital acts/week at baseline and 0.58 fewer at the last follow-up; and 1.9 coital act/week without condoms at baseline and 0.46 fewer at the last follow-up.
† Weighted average of 95.6% of participants with 8.2/month and 4.4% with <1/week.

Reported frequency of unprotected coital acts combined with estimated HIV prevalence in partners is sufficient to estimate women’s incidence from sex. To estimate partners’ prevalence, I use UNAID’s estimates of adult HIV prevalence for relevant countries and years (Annex Table 1). Assuming a rate of transmission of 0.001 (0.1%) per unprotected coital act with an HIV-positive partner, women in the six trials could be expected to contract HIV at rates of 0.2-1.2 (median 0.5) per 100 PYs (Table 2). These estimates may be too low (eg, because women misreported behavior). They may also be too high (eg, because PrEP protected women with sexual exposures).

With estimated rates of HIV incidence from reported sexual behavior, the estimated percent of infections from sex ranges from 3% (= 3/98) in CAPRISA 004 to 25% (= 17/68) in FEM-PrEP (median 13%)(see last two columns on the right in Table 2). Combining estimates from six trials, sex accounted for an estimated 89 infections or 11% of 779 observed infections, leaving an estimated 89% from bloodborne risks.

## Results: How did some RCTs succeed?

In six RCTs, incidence in control arms was 4.5-45 (median 11) times estimated incidence from sex (Table 2 and last two columns on the right in Table 3). If a majority of HIV comes from bloodborne risks, a successful intervention would have to reduce infections from bloodborne as well as sexual risks. This is reasonable for daily oral PrEP (cf: the Bangkok Tenofovir Study among IDUs reported HIV incidence 49% lower with oral TDF in the intervention arm vs. the placebo arm[26]).

**Table 3:**
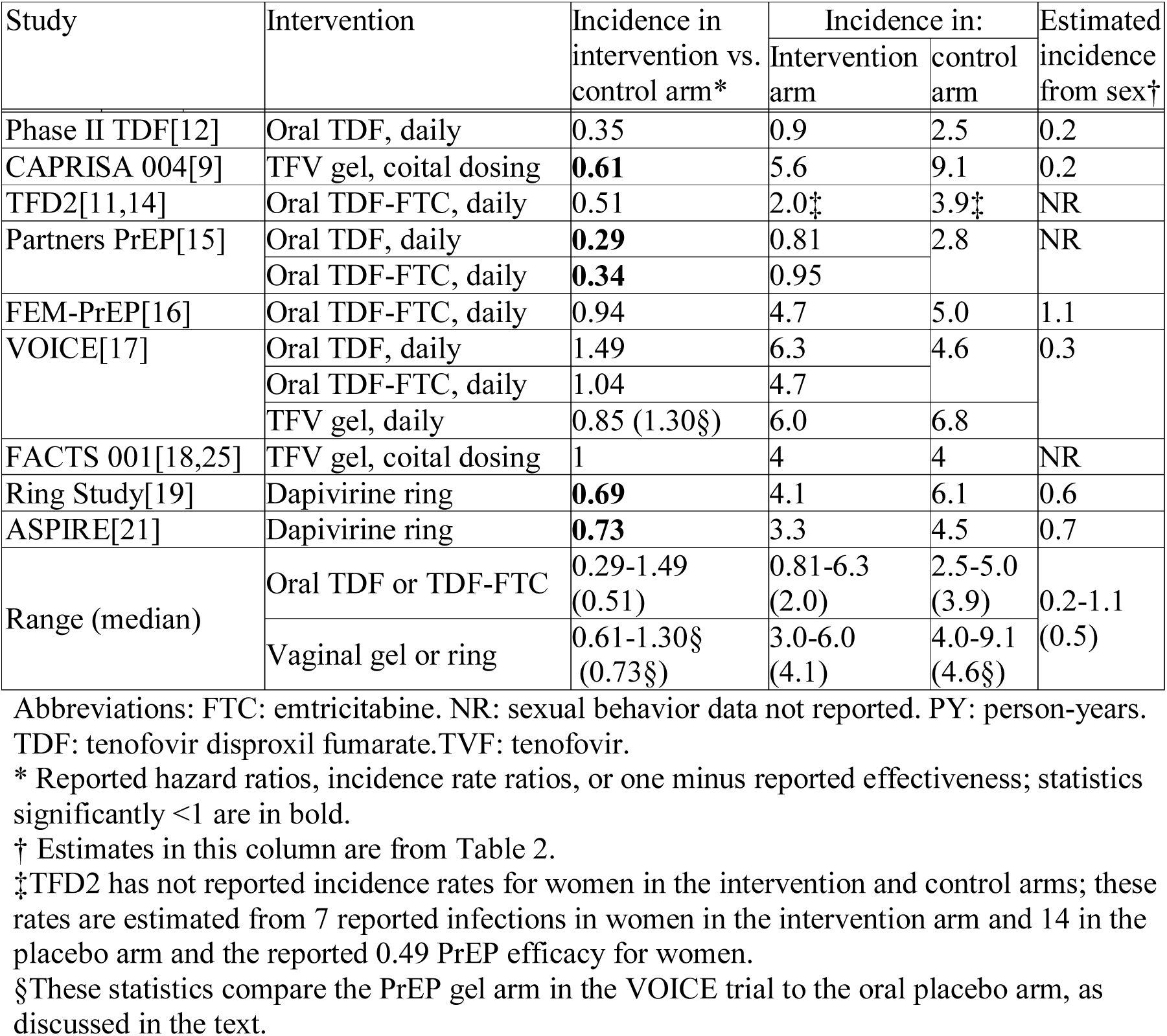
Incidence in intervention and control arms and estimated incidence from sex in nine RCTs

| Study | Intervention | Incidence in intervention vs. control arm* | Incidence in: |  | Estimated incidence from sex† |
| --- | --- | --- | --- | --- | --- |
|  |  |  | Intervention arm | control arm |  |
| Phase II TDF[12] | Oral TDF, daily | 0.35 | 0.9 | 2.5 | 0.2 |
| CAPRISA 004[9] | TFV gel, coital dosing | **0.61** | 5.6 | 9.1 | 0.2 |
| TFD2[11,14] | Oral TDF-FTC, daily | 0.51 | 2.0‡ | 3.9‡ | NR |
| Partners PrEP[15] | Oral TDF, daily | **0.29** | 0.81 | 2.8 | NR |
|  | Oral TDF-FTC, daily | **0.34** | 0.95 |  |  |
| FEM-PrEP[16] | Oral TDF-FTC, daily | 0.94 | 4.7 | 5.0 | 1.1 |
| VOICE[17] | Oral TDF, daily | 1.49 | 6.3 | 4.6 | 0.3 |
|  | Oral TDF-FTC, daily | 1.04 | 4.7 |  |  |
|  | TFV gel, daily | 0.85 (1.30§) | 6.0 | 6.8 |  |
| FACTS 001[18,25] | TFV gel, coital dosing | 1 | 4 | 4 | NR |
| Ring Study[19] | Dapivirine ring | **0.69** | 4.1 | 6.1 | 0.6 |
| ASPIRE[21] | Dapivirine ring | **0.73** | 3.3 | 4.5 | 0.7 |
| Range (median) | Oral TDF or TDF-FTC | 0.29-1.49 (0.51) | 0.81-6.3 (2.0) | 2.5-5.0 (3.9) | 0.2-1.1 (0.5) |
|  | Vaginal gel or ring | 0.61-1.30§ (0.73§) | 3.0-6.0 (4.1) | 4.0-9.1 (4.6§) |  |

Abbreviations: FTC: emtricitabine. NR: sexual behavior data not reported. PY: person-years. TDF: tenofovir disoproxil fumarate. TVF: tenofovir.
\* Reported hazard ratios, incidence rate ratios, or one minus reported effectiveness; statistics significantly <1 are in bold.
† Estimates in this column are from Table 2.
‡TFD2 has not reported incidence rates for women in the intervention and control arms; these rates are estimated from 7 reported infections in women in the intervention arm and 14 in the placebo arm and the reported 0.49 PrEP efficacy for women.
§These statistics compare the PrEP gel arm in the VOICE trial to the oral placebo arm, as discussed in the text.

Among the RCTs in this study, ratios of women’s incidence in intervention vs. control arms were ≤0.51 for only four interventions, all with oral PrEP (Table 3). Phase II TDF, the RCT for one of these interventions, disclosed reported sexual behavior sufficient to estimate HIV from sex: incidence in the control arm (2.5 per 100 PYs) was 12 times estimated incidence from sex (0.2 per 100 PYs). In this trial, PrEP may have succeeded by reducing both bloodborne and sexual transmission. Neither of the RCTs for the other three successful interventions (TDF2 and Partners PrEP) report women’s reported frequency of unprotected coital acts (in Partners PrEP among discordant couples, two frequencies would be required, for coital acts with primary partners and with other partners).

## Results: How did low adherence lead to poor results with oral PrEP?

Faced with disappointing results with oral PrEP among women in Africa (Table 3), the overwhelming response among study teams, reviewers, and supporting organizations has been to blame lack of success on low adherence.[22,24,27] In the three RCTs with the four successful interventions noted in the previous paragraph, women’s (or men’s and women’s) adherence to daily oral PrEP was 69%-80% (measured by counts of returned pills in one and plasma tests for drugs in two)(Tables 3 and 4). In the two RCTs with the three unsuccessful oral PrEP interventions (with ratios of incidence in intervention vs. control arms ranging from 0.94-1.49) women’s adherence to daily oral PrEP was only 29%-35% (measured by plasma tests for drugs)(Tables 3 and 4). Moreover, within trials with low adherence, HIV incidence was lower among women who used oral PrEP more consistently.[16,17] These observations agree with overwhelming evidence oral PrEP provides partial protection against HIV.

**Table 4:** Reported evidence on adherence vs. sexual risk in five trials of oral PrEP

| Study | Adherence (measure) | Reported evidence on women's awareness of sexual risk vs. adherence |
| --- | --- | --- |
| Phase II TDF[12] | ≤69% (pill count) | No report |
| TDF2[14] | 80% (drug in plasma) | No report. |
| Partners PrEP[15] | 80% (drug in plasma) | See text |
| FEM-PrEP[16] | ~35% (drug in plasma) | (a) The study team (p 420 in [16]) “hypothesize[d] that the women’s perception that they were at low risk for HIV infection may have contributed to the poor adherence”<br>(b) A sub-study reported some women stopped PrEP when they perceived no sex risk, ie, no partner[32]<br>(c) A sub-study reported good adherence was not associated with several measures of sexual risk (but measures in the analysis did not include not having sex)[33] |
| VOICE[17] | 29% (drug in plasma) | A sub-study reports women considered sexual risk when deciding whether or not to take study drugs[34,35]; eg, (pp 18-19 in [34]) “Some restarted using...when they felt an immediate risk from a promiscuous partner” and “...some women selectively used products when their partner was around...” |

Women’s reasons for low adherence are crucial to understanding what happened in these trials. If women used PrEP when they had sexual risks but not otherwise, and if all infections came from sex, then effectiveness should not depend on daily use (cf: an RCT of oral PrEP among MSM in France and Canada told men to use PrEP when they had sexual risks; incidence in the intervention arm was a significant 86% lower than in the placebo arm[28]). On the other hand, if most infections came from bloodborne risks, effectiveness would fall as women skipped daily doses, even if they scrupulously used PrEP for all sexual risks.

Three of the five trials of oral PrEP reported qualitative or quantitative evidence that women sometimes adhered or not according to sexual risks (Table 4). The best quantitative evidence relating adherence to women’s (and men’s) awareness of sexual risk comes from sub-studies linked to Partners PrEP:

- One sub-study reports significant adjusted odds ratios for low adherence (measured by electronic monitoring of pill bottle opening) of 1.30 for abstinence, 1.71 for sex with other partner (ie, not the HIV-positive primary partner) only with 100% condom use, and 2.48 for sex with other partner only with <100% condom use (vs. participants reporting sex with primary partner only and 100% condom use). Missing the study drug for ≥72 hours was significantly more frequent among participants reporting abstinence or sex with other partner only and <100% condom use (vs. participants reporting sex with primary partner only and 100% condom use).[29]
- Another sub-study with the same sample reports significant adjusted odds ratios for low adherence (measured by unannounced pill counts) of 4.2 for abstinence and 3.0 for sex with other partner + primary partner (vs. participants reporting sex with primary partner only and 100% condom use).[30]
- A sub-study using 60 consecutive daily phone surveys reports odds ratios for missing a daily PrEP dose of 3.29 for those with no sex in 60 days, 2.93 for those with 1-10 sex acts, and 1.96 for those with 11-20 sex acts vs. those with >20 sex acts in 60 days. The adjusted odds ratio for missing a daily PrEP dose was 1.87 for those not reporting sex that day vs. those reporting sex.[31]

## Results: evidence of failure and even harm with vaginal gels and rings

Three RCTs tested vaginal TFV gels (to be applied before and after coitus in two trials and daily in one); two tested Dapivirine vaginal rings, with instructions to use them continuously, inserting a new one every month. Reported protection was modest at best, with incidence in intervention vs. placebo arms ranging from 0.61-1 (median 0.73)(Table 3).

However, because both PrEP and placebo gels and rings may disturb the vagina in ways that could increase HIV risk,[36] testing vaginal PrEP against vaginal placebo may be misleading. For example, all three trials of PrEP gels used placebo gels with hydroxyethyl cellulose (HEC);[9,17,37] HEC has been associated with pro-inflammatory vaginal immune markers.[38] Among a sample of women in the CAPRISA 004 study, HIV incidence was associated with pro-inflammatory immune markers in plasma[39] and genital fluid.[40]

The VOICE trial – the only RCT offering a comparison between vaginal PrEP and oral placebo – found HIV incidence in both the TFV gel and placebo gel arms (6.0 and 6.8 per 100 PYs, respectively) to be greater than in the oral placebo arm (4.6 per 100 PYs). Comparing TFV gel to placebo gel, TFV gel appears to modestly reduce women’s risk, with an incident rate ratio of 0.85. However, a comparison of TFV gel vs. oral placebo (Table 3) suggests TFV gel increased women HIV incidence by 30% (= [6.0/4.6] - 1).

Although none of the other trials of gels or rings had an oral placebo arm, it is noteworthy that across all RCTs, median HIV incidence in arms with vaginal PrEP (4.1 per 100 PYs) was greater than median incidence in arms with oral placebos (3.9 per 100 PYs)(Table 3). Moreover, HIV incidence with vaginal PrEP was also high, ranging from 3.3-5.6 per 100 PYs in five intervention arms; these rates would bring women to 20% HIV prevalence in 2.6-6.1 years.

Although studies of vaginal PrEP reported evidence suggesting better protection with better adherence, the evidence is not as convincing as with oral PrEP. For example, both trials of Dapivirine rings reported only modestly lower incidence in intervention vs. control arms despite adherence exceeding 80% from tests of drug in returned rings and plasma. In FACTS 001 (p 2 in [41]) “[i]n) both the active and placebo arms of the trial, women who had high adherence as measured by returned applicators had lower rates of infection than women who had low adherence, …women who were adherent may have had lower risk because they were in some other ways different than non-adherent women.”

Finally, because PrEP in vaginal gels[9] and rings[21] gets into plasma, they could modestly impact bloodborne transmission,[42] though not as much as oral PrEP.

## Discussion

### Proportion of HIV incidence from bloodborne risks

The estimate in this study – that 11% of women’s HIV incidence across six RCTs came from sexual partners – challenges the dominant hypothesis (belief or assertion[43]) that most HIV in adults in Africa comes from sex. This challenge cannot be removed by tweaking the estimate; even arbitrarily tripling the estimate to 33% leaves two-thirds of infections from bloodborne risks.

Evidence from some other studies similarly shows a minority of HIV from sex among young women in southern Africa. For example, a 2012 survey of 1,698 young women aged ≥12 years (only 7% were ≥20 years old) in grades 8-12 in a high-prevalence sub-district in KwaZulu Natal, South Africa, found 54% (=56/104) of infections in self-reported virgins.[44] Using data from Demographic and Health Surveys in Lesotho, Malawi, and Swaziland, Eva Deuchert estimated 30% of HIV in unmarried women aged 15-19 years came from sex if all self-reported virgins were telling the truth; and that more than 55% of sexually active women would have had to misrepresent themselves as virgins for sex to account for more than half of observed HIV infections.[45]

Persistent evidence that heterosexual risks fall short of explaining most HIV in African adults challenges those who choose to focus on sex. For example, a prominent 2004 defense of the view that sex accounts for most HIV in Africa claims (pp 482, 484 in [46]) “epidemiological evidence indicates that sexual transmission continues to be by far the major mode of spread of HIV-1,” but belies that claim by dismissing conflicting evidence with the assertion “data on sexual behavior are notoriously imprecise …”

Estimates and evidence that non-sexual risks account for a majority of HIV in young women call attention to unreported evidence from the nine RCTs in this study. As of May 2017, no RCT has reported women’s HIV incidence during months with no coital act or 100% condom use vs. months with ≥1 unprotected coital act; these rates could be used to estimate proportions of infections from sex. Only two RCTs have reported any data at all on HIV incidence according to coitus during follow-up: (a) In Partners PrEP, men and women reporting any vs. no unprotected sex with (HIV-positive) primary partners had only moderately higher incidence, 2.4 to 1.5 per 100 PYs; the study did not say if they had other partners and did not sequence HIV to determine source.[15] (b) In ASPIRE, the rate of incidence in women reporting 0-1 sexual partners was 70% of the rate in women reporting ≥2 partners.[21] Also, but without disclosing data, the CAPRISA 004 study noted (p 995 in [39]) “neither coital frequency (protected or unprotected by condoms), nor the number of sexual partners…were strong predictors of HIV acquisition…”

### Implications for prevention: Warn women about bloodborne risks

In 2016 the UN General Assembly endorsed targets to reduce world HIV incidence from 1.9 million in 2015 to 500,000 in 2020.[47] If evidence-based estimates of HIV from sex among young women in southern and eastern Africa are anywhere near reality, prospects for success will depend on addressing not only sexual risks but also bloodborne risks.

Both UNAIDS and WHO have proposed strategies to reach the UN’s target. The five pillars of UNAIDS’ strategy are: sexual education and sexual health services, condoms, circumcising men, PrEP, and special attention to populations with specific risks (eg MSMs, IDUs).[48] UNAIDS’ strategy does not consider bloodborne risks other than IDU.

Along with interventions aimed at sexual risks and IDU, WHO’s strategy includes injection and blood safety, admonishing (p 33 in [49]) “safe medical injections and blood supplies, along with universal precautions, are central to a well-functioning health system.” To reduce bloodborne risks during medical injections, WHO recommends syringes with reuse prevention and safety features; aside from that supply side initiative, WHO’s strategy for bloodborne risks is short on details, with no mention of warning people about bloodborne risks and no mention of bloodborne risks in cosmetic services.

Some evidence suggests educating people to recognize and avoid bloodborne risks could have a big impact. During 2003-07, Demographic and Health Surveys in 16 African countries asked people what they could do to avoid getting AIDS; interviewers coded and recorded multiple answers. From 8.0% of women in Swaziland to 43.4% in Senegal provided answers coded as “avoid sharing razors/blades.” In the four countries where ≥30% of women mentioned avoiding sharing razors/blades (Ethiopia, Ghana, Rwanda, Senegal), women’s HIV prevalence ranged from 1.0% to 2.8% (median 2.4%). In the five countries where <15% of women provided such answers (Kenya, Lesotho, Swaziland, Tanzania, and Zimbabwe), from 6.4% to 26% (median 19.6%) of women were HIV-positive.[50]

The case for warning people about bloodborne risks is based not only on the potential for warnings to reduce HIV incidence but also on ethics. The World Medical Association’s Declaration of Lisbon[51] avers (principle 9): “Every person has the right to health education that will assist him/her in making informed choices about personal health and about the available health services…”

### Implications for prevention: Reconsider oral PrEP for women in Africa

Both UNAIDS and the World Health Organization (WHO) along with demonstration projects[52] propose extending oral PrEP in Africa to women in the general population who are at high risk (ie, in communities where HIV incidence in young women is >3 per 100 PYs).[53] If a majority of young women’s infections in such communities are from bloodborne risks in healthcare and cosmetic services, these recommendations should be reviewed: (a) to consider what if any role PrEP has to protect women from such risks; and (b) to reconsider PrEP as an option to protect women from revised lower estimates of HIV from sex.

## Acknowledgements

I thank Devon Brewer and Simon Collery and others for suggestions and advice. I received no funding for this study and have no conflict of interest with respect to this article.

---

**Appendix Table 1.** Estimated HIV prevalence in women’s sexual partners in six RCTs reporting women’s reported sexual behavior

| Study | Country | % of participants from the country | Study years | National adult HIV prevalence (average across study years)[1] | Estimated HIV prevalence in women's partners* |
| --- | --- | --- | --- | --- | --- |
| Phase II TDF[12] | CM | 43% (=371/859) | 2004 | 5.3% | 3.3% |
|  | GH | 42% (=361/859) | 2004-6 | 2.4% |  |
|  | NGA | 15% (=127/859) | 2004 | 3.9% |  |
| CAPRISA 004[9] | SA | 100% | 2007-10 | 19.5% | 19.5% |
| FEM-PrEP[16] | KY | 35% (=720/2056) | 2009-11 | 6.1% | 14.1% |
|  | SA | 62% (=1,279/2,056) | 2009-11 | 19.0% |  |
|  | TZ | 3% (=57/2,056) | 2009-11 | 5.6% |  |
| VOICE[17] | SA | 81% (=4,077/5,029) | 2009-12 | 18.9% | 17.7% |
|  | UG | 6% (=322/5,029) | 2009-12 | 7.1% |  |
|  | ZI | 13% (=630/5,029) | 2009-12 | 15.4% |  |
| Ring Study[19,20] | SA | 90% (=1,762/1,959) | 2012-15 | 19.0% | 17.8% |
|  | UG | 10% (=197/1959) | 2012-15 | 7.2% |  |
| ASPIRE[21] | MW | 10% (=272/2,629) | 2012-15 | 9.7% | 15.9% |
|  | SA | 54% (=1,426/2,629) | 2012-15 | 19.0% |  |
|  | UG | 10% (=253/2,629) | 2012-15 | 7.2% |  |
|  | ZI | 26% (=678/2,629) | 2012-15 | 15.0% |  |

Abbreviations: BT: Botswana. CM: Cameroon. GH: Ghana. KY: Kenya. MW: Malawi. NGA: Nigeria; SA: South Africa. TZ: Tanzania. UG: Uganda. ZI: Zimbabwe.
\* Average HIV prevalence in adults weighted by countries and years.

